# Energy demand and the context-dependent effects of genetic interactions

**DOI:** 10.1101/209510

**Authors:** Luke A. Hoekstra, Cole R. Julick, Katelyn M. Mika, Kristi L. Montooth

## Abstract

Genetic effects are often context dependent, with the same genotype differentially affecting phenotypes across environments, life stages, and sexes. We used an environmental manipulation designed to increase energy demand during development to investigate energy demand as a general physiological explanation for context-dependent effects of mutations, particularly for those mutations that affect metabolism. We found that increasing the photoperiod during which *Drosophila* larvae are active during development phenocopies a temperature-dependent developmental delay in a mitochondrial-nuclear genotype with disrupted metabolism. This result indicates that the context-dependent fitness effects of this genotype are not specific to the effects of temperature and may generally result from variation in energy demand. The effects of this genotype also differ across life stages and between the sexes. The mitochondrial-nuclear genetic interaction disrupts metabolic rate in growing larvae, but not in adults, and compromises female, but not male, reproductive fitness. These patterns are consistent with a model where context-dependent genotype-phenotype relationships may generally arise from differences in energy demand experienced by individuals across environments, life stages, and sexes.

**IMPACT SUMMARY:** Genetic effects on traits are often context dependent, such that a genotype that improves fitness under one context may have no effect or even a deleterious effect in another context. The external environment is a common context that affects the degree to which a genotype determines a phenotype, but the internal environment of an organism (e.g., its genetic background, sex or life stage) also provides an important context that may modify the phenotypic expression of a genotype. Here we combine new data on the phenotypic effects of a well-characterized genetic interaction between the mitochondrial and nuclear genomes of the fruit fly *Drosophila* with prior observations to support a model of energy demand as a general explanation for context-dependent genetic effects, particularly for mutations that affect metabolism. We show that the magnitude of fitness effects of this genetic interaction correlates positively with the degree of energy demand among developmental treatments that accelerate growth rate, across developmental stages that differ in the cost of growth, and between sexes with potentially different costs of reproduction. These internal and external contexts create variable demands on energy metabolism that will impact the efficacy of natural selection acting on metabolic mutations in populations.

## INTRODUCTION

Environment, development and physiological state can all modify the phenotypic expression of genetic variation (e.g. Hartman *et al.* 2001; Raj *et al.* 2010). Because natural selection acts upon the subset of expressed genetic variation that affects fitness, the fate of new mutations may depend on the landscape of genetic backgrounds and environments experienced by a population (e.g. Remold and Lenski 2004, Chandler *et al.* 2013, Lachance *et al.* 2013, Wang *et al.* 2013, Kammenga *et al.* 2017). Routine variation in the internal – genetic, developmental, physiological – or external environment can challenge the capacity of individuals to maintain homeostasis, and this can magnify deleterious mutational effects (e.g. Kondrashov and Houle 1994, Hoekstra *et al.* 2013). Conversely, favorable environments can mask potential genotype-phenotype relationships (Harshman and Zera 2007, Agrawal *et al.* 2010, Hoekstra *et al.* 2013). If the relationship between genotype and fitness is generally conditional on internal or external environmental factors (i.e., is context dependent), then elucidating general principles underlying genotype-phenotype-environment interactions is critical for understanding evolutionary processes such as the maintenance of genetic variation for life-history traits (Roff and Fairbairn 2007, Van Dyken and Wade 2010, Mackay 2013).

If context-dependent genetic effects mediate phenotypic tradeoffs – often manifest as negative phenotypic trait correlations – then dissecting the underlying physiology can provide mechanistic explanations of phenotypic correlations that better enable predictions regarding the performance of particular genotypes in particular environments (Harshman and Zera 2007, Flatt and Heyland 2011). Many phenotypic tradeoffs likely result from the differential allocation of finite resources to growth, survival, and reproduction (Van Noordwijk and Dejong 1986, Roff 2002). Traits such as growth rate and gamete production demand sufficient energy production supplied by metabolic processes, but the rate of metabolism itself is subject to homeostatic regulation that can influence energy allocation and obscure trait correlations (Clarke and Fraser 2004, Harshmann and Zera 2007, Leopold and Perrimon 2007). While many good examples of the importance of environmental context for tradeoffs exist (reviewed in Asplen *et al.* 2012), understanding the genetic architecture underlying tradeoffs and the physiological mechanisms mediating them lags behind (Roff and Fairbairn 2007).

To begin to fill this gap in our understanding, we combined well-characterized mitochondrial-nuclear (hereinafter mito-nuclear) genotypes that affect metabolism, physiology and fitness in Drosophilid flies(Hoekstra *et al.* 2013, Meiklejohn *et al.* 2013, Holmbeck *et al.* 2015, Zhang *et al.* 2017) with an environmental perturbation designed to manipulate energy demand. Phenotypic effects of variation in the mitochondrial genome (mtDNA) often depend upon variation in the nuclear genome due to the functional interactions between gene products from these two genomes (reviewed in Burton and Barreto 2012). In ectotherms, the phenotypic effects of these mito-nuclear genetic interactions frequently depend upon temperature (Dowling *et al.* 2007a; Arnqvist *et al.* 2010; Hoekstra *et al.* 2013; Paliwal *et al.* 2014). For example, cool development temperatures masked the deleterious effects of a mito-nuclear incompatibility in *Drosophila*, while warmer temperatures generated inefficiencies in larval metabolism that magnified the deleterious effects of the incompatibility on development rate (Hoekstra *et al.* 2013). We proposed that this was due to the accelerating effect of temperature in speeding up development and increasing demand on energetic processes during more rapid larval growth. Here we show that manipulating the developmental photoperiod to accelerate growth and putatively increase energy demand (Kyriacou *et al.* 1990, Paranjpe *et al.* 2005) generates similar context-dependent effects of this genotype on development that are independent of temperature. This suggests that energy demand may be a general physiological explanation for why some environments expose, while others mask, genetic effects.

Energy demand may also provide a general explanation for why genetic effects vary across life stages and between sexes. The substantial metabolic cost of growth (Parry 1983, Glazier 2005) may cause the energy budget of developing organisms to be more constrained than that of adults. This may be particularly true for holometabolous insects that experience exponential growth during development before reaching a relatively static adult size (Church and Robertson, 1966). After the cessation of growth, adults of different sexes may partition energy in different ways due to the differential costs of reproduction (Bateman 1984; Hayward and Gillooly 2011), potentially generating sex-specific effects of mutations. Here we present patterns of context-dependent effects of a mito-nuclear incompatibility that are consistent with a model where internal and external environments that cause energy demand to exceed supply may generally expose mutational effects on phenotypes. This has important consequences for the efficacy of natural selection acting on mutations in populations, and particularly for those mutations that impact metabolism.

## METHODS

### Drosophila *genotypes*

We used mito-nuclear genotypes that precisely pair mtDNAs from *Drosophila melanogaster* and *D. simulans* with nuclear genomes from wild-type *D. melanogaster* (Montooth *et al.* 2010). Pairing the *D. simulans* mtDNA from the *simw^501^* strain with two *D. melanogaster* nuclear genomes reveals a strong mito-nuclear epistatic interaction for fitness. The *D. simulans simw^501^* mtDNA is phenotypically wild type when combined with the *D. melanogaster AutW132* nuclear genome (hereinafter *Aut*), but is incompatible with the *D. melanogaster Oregon-R* (hereinafter *OreR*) nuclear genome, resulting in a significant increase in development time and decrease in fitness of the (*simw^501^*);*OreR* (mtDNA);nuclear genotype (Montooth *et al.* 2010, Meiklejohn *et al.* 2013). The molecular genetic basis of this interaction is an incompatible pairing between a single nucleotide polymorphism (SNP) in the *simw^501^* mitochondrial-encoded tRNA^Tyr^ and a naturally segregating amino acid polymorphism present in the *OreR* nuclear-encoded mitochondrial tyrosyl-tRNA synthetase gene, *Aatm*. These mutations act epistatically to decrease OXPHOS activity, as predicted by the critical role that these genes play in mitochondrial protein translation (Meiklejohn *et al.* 2013). The four genotypes that we use here – (*ore*);*Aut*, (*ore*);*OreR*, (*simw^501^*);*Aut*, and (*simw^501^*);*OreR* – provide a well-characterized model of epistasis between naturally occurring polymorphisms that affects energy metabolism and fitness, allowing us to test how internal and external environment influences the phenotypic expression of genetic interactions (Table 1). Fly cultures were maintained on Bloomington *Drosophila* Stock Center media with a 12:12h light:dark cycle, unless otherwise indicated.

**Table 1.**
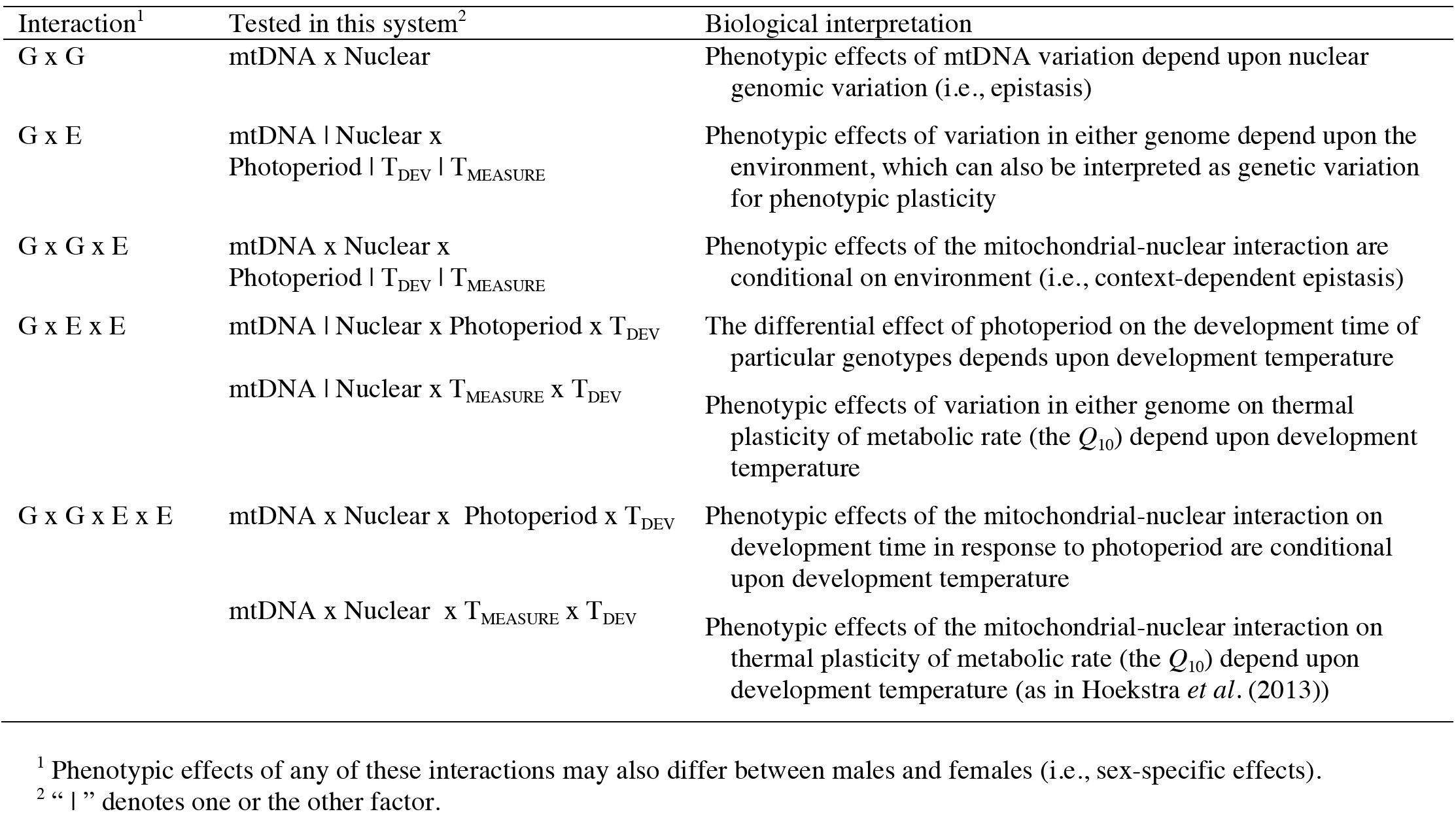
Biological interpretation of context-dependent genetic effects in this study system

### Manipulating developmental photoperiod

We tested the specific prediction that an arrhythmic photoperiod (24:0h L:D) would accelerate growth rate in wild-type genotypes, but that the increased energy demand of accelerated growth would induce a developmental delay of the incompatible (*simw^501^*);*OreR* genotype at 16°C – a temperature where this genotype has a wild-type development time (Hoekstra et al. 2013). We also tested whether arrhythmic photoperiods at 22aC would phenocopy the developmental delay caused by (*simw^501^*);*OreR* at higher developmental temperatures (Hoekstra *et al.* 2013). We quantified the effect of extended day length on development time using four different combinations of temperature and light:dark cycle (16°C, 12:12h; 16°C, 24:0h; 22°C, 12:12h; 22°C, 24:0h). For each genotype, replicate pools of fifty 0-12 hour old eggs were collected into fresh food vials and randomly assigned to one of the four developmental treatments. We scored the number of new pupae and new adults to eclose once per day at 16°C and twice per day at 22°C for approximately 20 vials of each genotype under each developmental treatment. The fixed effects of genotype and photoperiod on development time within each temperature were tested using mixed-model analysis of variance (ANOVA) models that were fit using restricted maximum likelihood and included rearing vial as a random factor. All analyses were performed in the R statistical package (R Core Team 2013).

### Adult mass and metabolic rate

Incompatible (*simw^501^*);*OreR* larvae have inefficient larval metabolism, manifest as higher metabolic rates and longer development times at 25°C, but develop and respire at a normal pace at 16°C (Hoekstra *et al.* 2013). To test whether this mito-nuclear incompatibility also affects adult metabolism, we reared all four genotypes from egg to adult with controlled densities at 16°C or 25°C and measured mass and metabolic rate of 3-6 day old adults. At 48 hrs. post-pupal eclosion, adult flies were lightly anaesthetized with CO_2_, sexed, and sorted into groups of ten flies. The wet mass of each group of ten flies was recorded to the nearest μg and adults were allowed to recover in fresh, yeasted food vials for at least 24hrs. Mass was log-transformed to improve normality and genetic effects on mass were tested using ANOVA and Tukey’s post-hoc contrasts corrected for the number of multiple tests.

We used flow-through respirometry to estimate routine metabolic rate (hereinafter, metabolic rate) as the volume of CO_2_ (VCO_2_) produced by groups of ten female or male flies of the same genotype that were confined to a small, dark space to minimize activity. VCO_2_ is a good proxy for metabolic rate in insects like *D. melanogaster* that largely use carbohydrates for respiration and have a respiratory quotient of approximately one (Chadwick 1947). We measured at least ten biological replicates of each combination of genotype, sex, development temperature (T_DEV_ = 16°C and 25°C), and measurement temperature (T_MEASURE_ = 16°C and 25°C) using offspring collected from multiple cultures and multiple parental generations in order to average
across micro-environmental effects. Metabolic rates were measured between 11:00 am and 7:00 pm, and genotypes were distributed across this timeframe and across respirometry chambers using a random, balanced design. All measurements were made in a Peltier-controlled thermal cabinet (Tritech Research, Inc.), and measurement temperature was monitored using a thermocouple meter wired into an empty respirometry chamber.

For flow-through measurement of adult VCO_2_ we pushed CO_2_-free air through glass respirometry chambers containing flies at a rate of 100 ml min^-1^. Air that leaves the chamber carries CO_2_ produced by the flies, as well as water. The water vapor was removed from the airstream using magnesium perchlorate, and the CO_2_ in the airstream was measured using a Licor 7000 infrared CO_2_ detector (Licor, Lincoln, NE, USA). We used the RM8 Intelligent Multiplexer to switch the airstream sequentially through five respiratory chambers (Sable Systems International, Las Vegas, NV, USA), one of which serves as an empty baseline chamber. Each experimental run measured the VCO_2_ of four pools of ten adult flies, each sampled twice for ten minutes during a 100-minute period. There is no death from this treatment.

Baseline CO_2_ values were recorded before and after each sample and used to drift-correct CO_2_-tracings using the two-endpoint automatic method in Expedata, version 1.1.15 (Sable Systems International, Las Vegas NV). Raw CO_2_ values were converted from parts per million to μL hr^-1^ (VCO_2_) and then log-transformed to improve normality and homoscedasticity. To allow metabolic rate to acclimate in response to temperature shifts (e.g., T_DEV_ = 16°C and T_MEASURE_ = 25°C), we used the second recording of each respirometry run to estimate metabolic rate such that flies were acclimated for 50-80min.

Because there is measurement error in adult body mass, we estimated the scaling relationship between mass and VCO_2_ using Type II Model regression implemented with smatR, version 3.4.3 (Warton *et al.* 2006). When justified by a homogeneity of slopes test for the log-log relationship between mass and VCO_2_, we fit a common slope to all genotypes and tested for shifts along the common x axis (i.e., differences in mass) and for shifts in elevation (i.e., differences in mass-specific metabolic rate) among genotypes. Across genotypes within each of the four T_MEASURE_ x Sex combinations, we were able to fit a common slope and test for genotype differences in mass and in mass-specific metabolic rate. We could then correct for the effect of mass on metabolic rate by taking the residuals of each of these regressions and adding back the grand mean of all fitted values to provide meaningful scale. We refer to these values as mass-corrected metabolic rates (MCMR). We tested for the fixed effects of T_DEV_, mtDNA, nuclear genotype, and all possible interactions on MCMR using analysis of variance (ANOVA).

Respirometry chambers were housed inside infrared activity detectors (AD-2, Sable Systems International, Las Vegas NV), providing a simultaneous measurement of activity. We summarized the activity data by taking the median absolute difference of activity across each seven-minute metabolic rate measurement. This measure of activity neither significantly affected metabolic rate nor interacted with any genetic effects and was not included in our final statistical models.

### Adult reproductive traits

Incompatible (*simw^501^*);*OreR* females have significantly reduced fecundity, measured as the number of eggs laid over 10 days (Meiklejohn *et al.* 2013). To test whether males of this genotype also suffer a decrease in reproductive fitness, we measured one aspect of male fertility – the number of offspring sired by an individual male mated to virgin females of a control wild-type genotype, *Canton-S*. Thirty males were assayed for fertility across two experimental blocks that spanned multiple parental generations and used slightly different female genotypes. Both female genotypes were *Canton-S*, but in the first block the strain carried the *cn,bw* eye mutation. Block was included as a fixed factor in the analysis. However, rank orders of genotypes for the number of females fertilized and the number of offspring sired were similar between blocks.

Males were given 48-hours to mate with three virgin females. After 48-hours, we placed each female into a separate vial to lay fertilized eggs for an additional week. Progeny emerging from each vial were counted every other day until all progeny were counted. Males that sired fewer than 25 offspring were removed from the analysis, which resulted in a sample size of 29 males per genotype, except for (*simw^501^*);*Aut* for which n = 28 males. By placing females in separate vials after they were housed with the focal male, we could infer how many females were mated by each male, with the caveat that some females may have laid eggs in the first vial, but not in their individual vials. The median number of females that produced offspring in their vial per male was 3 (mean = 2.64) and this was not affected by genotype (*P* > 0.15 for genotype and all interactions). We estimated fertility as the total progeny sired by each male divided by the
number of females with whom that male produced progeny. We tested for fixed effects of mtDNA, nuclear genotype, experimental block and the interaction between these factors using ANOVA.

## RESULTS

### An arrhythmic photoperiod phenocopies the temperature-dependent developmental delay of a mito-nuclear incompatibility

The developmental delay of incompatible *(simw^501^);OreR* larvae is strongly mediated by temperature, with warmer temperatures exacerbating and cooler temperatures masking the delay (Hoekstra *et al.* 2013). *Drosophila* development rate depends on variation in the rhythmicity of the photoperiod (e.g. Kyriacou *et al.* 1990, Paranjpe *et al.* 2005). We tested whether an arrhythmic, continuous-light photoperiod (24:0h L:D) that typically accelerates larval-to-adult development (Paranjpe *et al.* 2005) could phenocopy this developmental delay in a manner similar to the accelerating effect of increased temperature. The accelerating effect of arrhythmic photoperiod influenced the severity of the *(simw^501^);OreR* developmental delay in a pattern remarkably similar to the effect of increased development temperature (Figure 1, Supplemental Tables S1 & S2). The incompatible *(simw^501^);OreR* genotype developed at the same pace as other genotypes when reared at 16°C with a fluctuating photoperiod (12:12h L:D) (mtDNA x Nuclear: *F_1, 80_* = 0.38, *P* = 0.5413). However, the incompatible *(simw^501^);OreR* genotype experienced a significant developmental delay relative to other genotypes when developed at 16°C with constant light (24:0h L:D) (mtDNA x Nuclear: *F_1, 79_* = 30.13, *P* <0.0001). At 22°C, where the incompatible genotype normally experiences a significant developmental delay with a fluctuating photoperiod, the constant light photoperiod magnified the developmental delay (Photoperiod x mtDNA x Nuclear: *F_1, 153_* = 299.07, *P* <0.0001). This was a particularly striking effect; constant light accelerated development of compatible mito-nuclear genotypes by ~ 2 days at 22°C, which is comparable to development time at 25°C. In contrast, constant light at 22°C significantly slowed development of the incompatible *(simw^501^);OreR* genotype, resulting in an ~ 4-day developmental delay between *(simw^501^);OreR* and compatible genotypes (Figure 1). Within each developmental temperature, the magnitude of the effect of the mito-nuclear genetic interaction on development time was conditional on photoperiod (G x G x E, Table 1; *P*<0.0001 for both developmental temperatures, Supplemental Table S1). However, there was no evidence that the highest-order interaction between development temperature, photoperiod, mtDNA and nuclear genotype affected development time (G x G x E x E, Table 1; *F_1,312_* = 0.41, *P* =0.5244). In other words, the effects of development temperature and photoperiod to delay development specifically in the incompatible genotype were independent.

**Figure 1.**
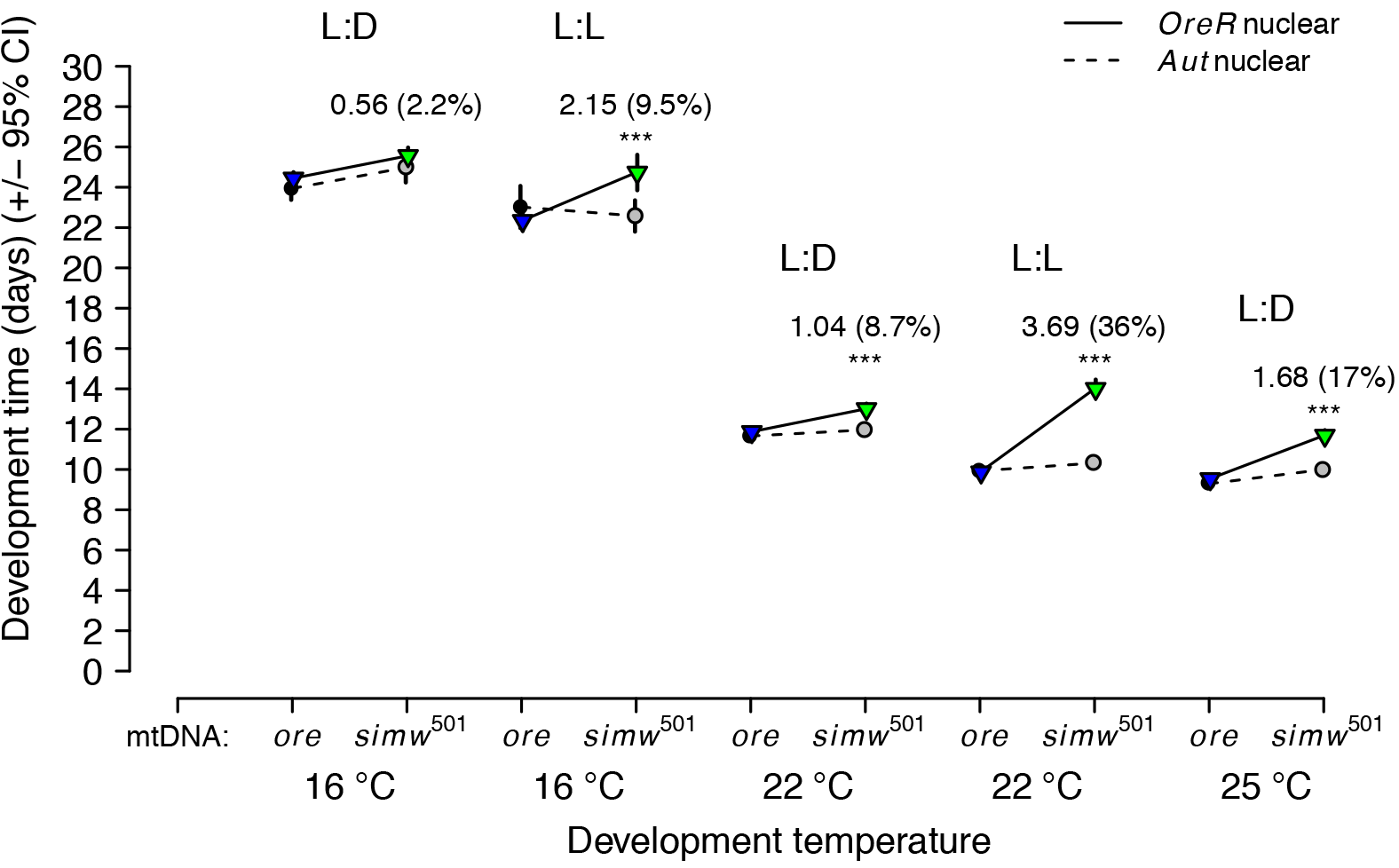
An arrhythmic photoperiod phenocopies the effects of increasing temperature on a mito-nuclear incompatibility that extends development. An arrhythmic, constant light photoperiod (L:L) accelerates development in control genotypes relative to a rhythmic photoperiod (L:D), but delays development in the mito-nuclear incompatible genotype (*simw^501^*);*OreR* (Supplemental Table S1). For comparison, the developmental delay of (*simw^501^*);*OreR* larvae relative to control genotypes under constant light at 22°C is greater than the delay observed at 25°C under fluctuating light (L:D) (25°C data from Hoekstra *et al.* (2013)). Asterisks denote a significant effect of the mtDNA x nuclear genetic interaction within each temperature-photoperiod combination at the level of *P* < 0.0001 (Supplemental Table S2), with the associated mean days delayed of (*simw501*);*OreR* relative to (*simw^501^*);*Aut* and the percent increase in development time in parenthesis.

### Adults with a mito-nuclear incompatibility achieve similar masses as compatible genotypes

Gene-environment (G x E) effects on adult mass were dominated by interactions between nuclear genotype and the known effects of both development temperature and sex on mass (Supplemental Table S3). Relative to these large effects, there was a small, but statistically significant effect of the mito-nuclear interaction on mass (mtDNA x nuclear: *F_1,317_* = 6.078, *P* = 0.0142). Rather than eclosing as smaller adults, (*simw^501^*);*OreR* adults were slightly larger than (*ore*);*OreR* adults (*P_Tukey_* = 0.04). However, the magnitude of the effect was small (+ 0.04 mg/10 flies) and the mtDNA x nuclear interaction was only statistically significant for females raised at 16°C (Supplemental Table S4). This small but consistent effect may be caused by smaller (*simw^501^*);*OreR* larvae not surviving development as well as smaller individuals of other genotypes, as this genotype has reduced egg-to-pupal survival (Hoekstra *et al.* 2013). In summary, adults with a mito-nuclear incompatibility that survive development achieve body masses that are similar to or greater than compatible genotypes, potentially as a consequence of their extended time spent as feeding larvae.

### In contrast to larvae, adult metabolic rate is more robust to mito-nuclear interactions

The scaling of metabolic rate as a function of mass can be characterized by the slope of the relationship between *ln*(metabolic rate) and *ln*(mass) (i.e., the mass-scaling exponent). The mass-scaling exponent differed slightly, but significantly between sexes and between measurement temperatures for adults in this study (*P* < 0.0001 for both Sex x Mass and T_MEASURE_ x Mass interactions). However, within each combination of sex and T_MEASURE_, there was no evidence that genotype or development temperature affected the mass-scaling exponent (*P* > 0.05 for both). Thus, we fit common slopes to the metabolic rate data for all genotypes within sex- T_MEASURE_ combination to test for effects of mito-nuclear genotype (Supplemental Figure S1 and Table S5). Within each sex-T_MEASURE_ combination, the range of masses for all genotypes were largely overlapping and there was no evidence for differences in mass among genotypes (Supplemental Figure S1 and Table S5).

In contrast to the larval life stage where *(simw^501^);OreR* larvae have significantly elevated 25lC metabolic rates (Hoekstra et al. 2013), there was no evidence of elevated metabolic rate in the incompatible *(simw^501^);OreR* adults measured at any temperature. The only significant effect of the mito-nuclear genotype on adult metabolic rate was a lower *(simw^501^);OreR* metabolic rate relative to compatible genotypes at 16°C. However, once again, this effect was small and only in females (Figure 2A, Supplemental Figure S1 and Table S5). Male metabolic rate was unaffected by mito-nuclear genotype (Supplemental Figure S1 and Table S5). Significant effects of the nuclear genome at 25°C (Supplemental Table S5) and in our prior work (Hoekstra & Montooth 2013, Hoekstra et al. 2013, Greenlee *et al.* 2014) demonstrate the sensitivity of this method to detect both genetic and environmental effects on metabolic rate.

**Figure 2.**
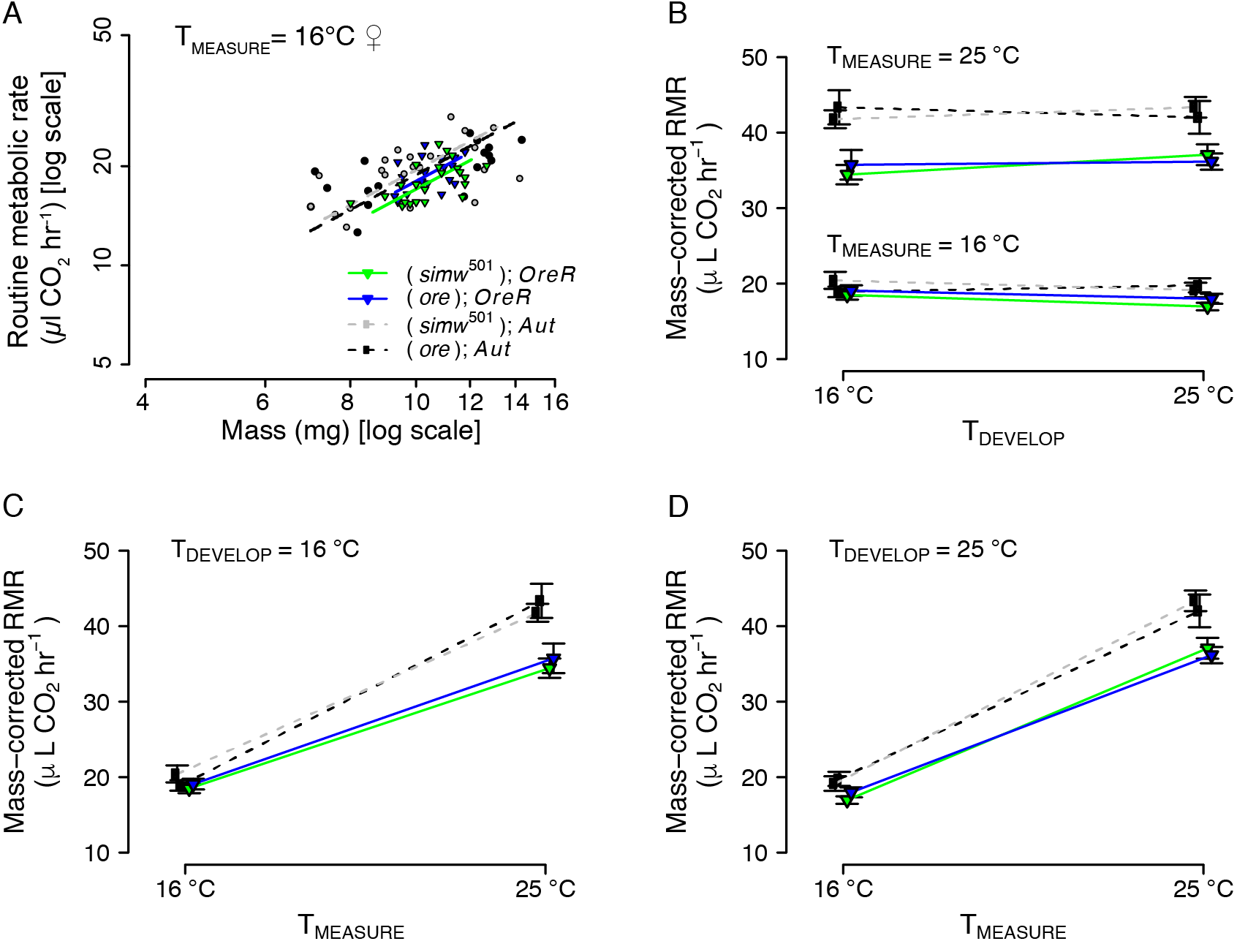
Female adult metabolic rate is robust to mito-nuclear genetic effects. A. The only mito-nuclear genetic effect was a small, but significant decrease in 16°C routine metabolic rate (RMR) in incompatible (*simw^501^*);*OreR* females (*P* < 0.05, Supplemental Table S5). B. Developmental reaction norms show robust thermal acclimation of female mass-corrected RMR at both measurement temperatures. C & D. Thermal reaction norms show that the *Q_10_* for female mass-corrected RMR is similar under both developmental temperatures, and that incompatible (*simw^501^*);*OreR* females have similar metabolic plasticity as their nuclear genotypic control (*ore*);*OreR*. Patterns were similar for adult male metabolic rate (Supplemental Figure S2). The mtDNA x nuclear interaction did not affect mass-corrected RMR of males or females at either measurement temperature (*P* > 0.28; Supplemental Table S6). Error bars are +/− 1 SEM.

Mass-corrected metabolic rates allow for comparisons of genotypes across development and measurement temperatures (i.e., metabolic plasticity) (Figure 2B-D, Supplemental Figure S2, Supplemental Tables S6 & S7). This analysis revealed that the lower 16°C metabolic rates in (*simw^501^*);*OreR* females were largely the consequence of lower metabolic rates of females developed at 25°C and measured at 16°C. However, this difference was not statistically significant and supports generally weak effects of this genetic interaction on adult, relative to larval, metabolic rates. There was no evidence that the mito-nuclear interaction affected adult male MCMR within any combination of development or measurement temperatures (Supplemental Fig S2 and Table S6). In contrast to larvae, where (*simw^501^*);*OreR* have compromised thermal plasticity of metabolic rate (i.e., the *Q_10_* for metabolic rate), we found no evidence that adults of this genotype differ in their *Q_10_* for metabolic rate (T_MEASURE_ x mtDNA x nuclear, *P* > 0.65 for both sexes) and this was independent of development temperature (T_MEASURE_ x T_DEV_ x mtDNA x nuclear, *P* > 0.50 for both sexes)(Supplemental Table S7). Thus, relative to larvae, both adult metabolic rate and metabolic plasticity are more robust to the effects of this mito-nuclear genetic incompatibility.

### Mito-nuclear effects on reproductive fitness are stronger in females

Patterns of male fertility and female fecundity provide further evidence that the effects of the mito-nuclear incompatibility are sex specific. The mito-nuclear interaction did not compromise the number of offspring sired by males (mtDNA x nuclear: *F_1, 107_* = 0.185, *P* = 0.668)(Figure 3A, Supplemental Table S8). This is in contrast to strong mito-nuclear effects on female fecundity, with females of the *(simw^501^);OreR* genotype producing ~50% fewer eggs than (*ore*);*OreR* females (Figure 3B, Supplemental Table S8)(Meiklejohn *et al.* 2013). Thus, while both sexes generally maintain metabolic rate independent of mito-nuclear genotype, *(simw^501^);OreR* females appear to do so at the cost of egg production.

**Figure 3.**
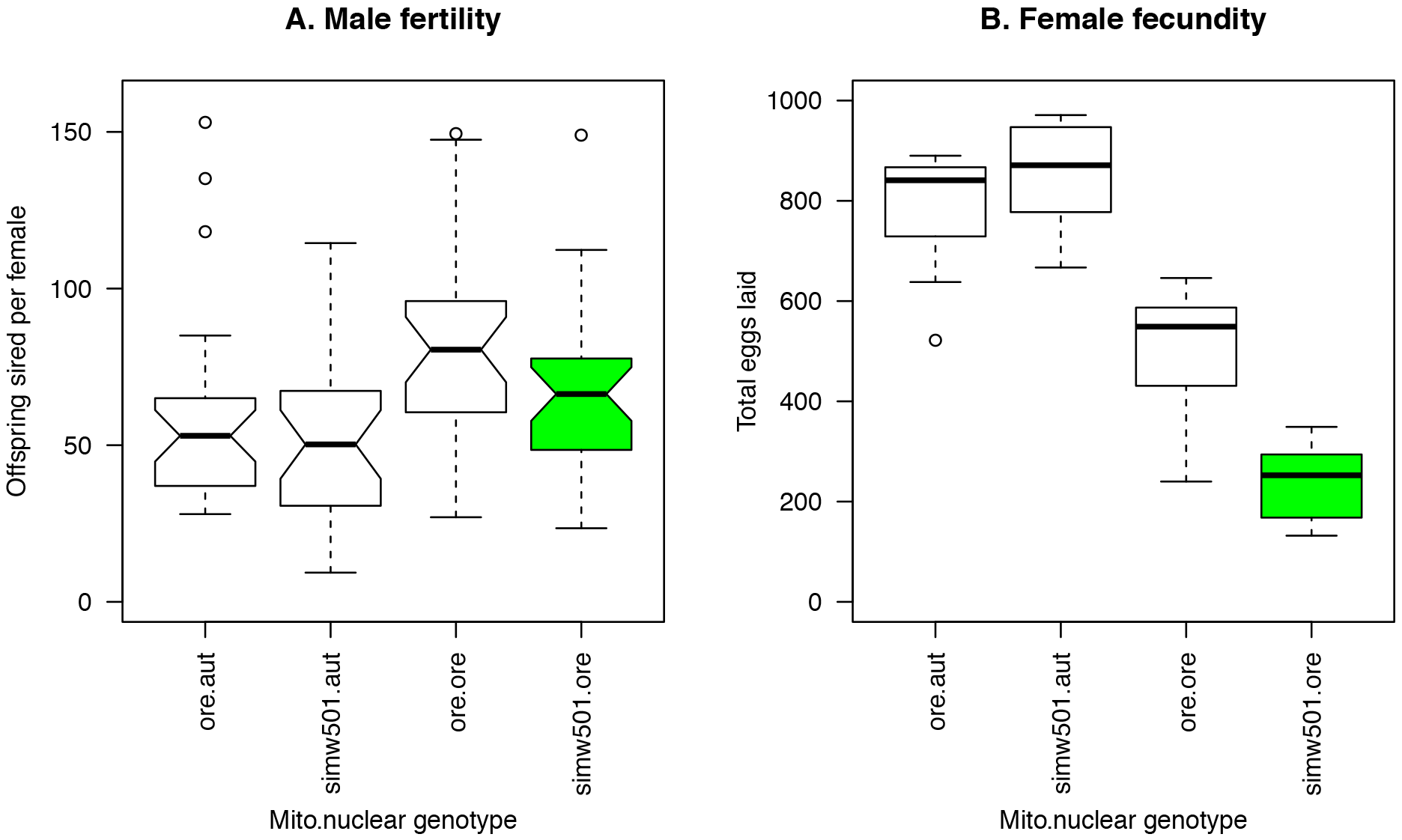
Mito-nuclear effects on reproductive phenotypes are stronger in females. A. Male fertility was not significantly affected by the mito-nuclear incompatibility (mtDNA x nuclear: *F_1, 107_* = 0.185, *P* = 0.668). B. This was in contrast to strong effects of mito-nuclear genotype on female fecundity (mtDNA x nuclear: *F_1, 31_* = 16.976, *P* = 0.0003), with females of the incompatible *(simw^501^);OreR* genotype laying on average 48% the number of eggs produced by *(ore);OreR* females. Female fecundity data from Meiklejohn *et al.* (2013) are total eggs produced over 10 days by individual females for n = 7-10 females per genotype. Whiskers on boxplots represent interquartile ranges. Green denotes the mitochondrial-nuclear incompatible genotype, as in other figures.

## DISCUSSION

Previously, we found that a synergistic, epistatic interaction between polymorphisms in the mt- tRNA^Tyr^ and the nuclear-encoded, mitochondrially targeted tRNA-synthetase for this mt-tRNA decreases OXPHOS activity and negatively impacts a number of life-history traits in a temperature-dependent manner in *Drosophila* (Montooth *et al.* 2010, Meiklejohn *et al.* 2013, Hoekstra *et al.* 2013, Zhang *et al.* 2017). Here we show that the phenotypic expression of this genetic interaction is generally environment-dependent and depends on intrinsic factors such as life stage and sex in a manner consistent with greater energy demands revealing fitness effects of mutations that compromise metabolism.

### The physiological basis of environment-dependent, mito-nuclear effects

Many organismal traits, particularly in ectotherms, are temperature dependent, including the universal, but mechanistically not well understood, unimodal thermal performance curve for metabolic rate (Angiletta 2009, Schulte 2015, DeLong *et al.* 2017). Given this relationship between temperature and metabolic processes, the thermal environment is likely an ecologically relevant and critical determinant of the relationship between genotype and phenotype for traits that depend on metabolic function. Phenotypic effects of cytonuclear interactions, including the mito-nuclear incompatibility described here, can be temperature sensitive (Arnquvist *et al.* 2010, Dowling *et al.* 2007a, Hoekstra *et al.* 2013). The temperature dependence of these mito-nuclear genetic effects could result from direct thermodynamic effects on the physical interaction between the mutations, e.g., between mutations in the mt-tRNA^Tyr^ and its tRNA synthetase. Alternatively, the temperature dependence may arise indirectly, as metabolic rate and development rate increase with temperature and place greater demand on the energetic products of mito-nuclear interactions. This latter, energy-dependent explanation is also consistent with the observations that mitochondrial genetic effects can be temperature sensitive (Pichaud *et al.* 2013) and both mitochondrial and mito-nuclear effects can be diet sensitive (Zhu *et al.* 2014, Ballard and Youngson 2015, Mossman *et al.* 2016). Here we found that manipulating the photoperiod to accelerate growth rate independent of temperature produced patterns of context-dependent mito- nuclear genetic effects strikingly similar to those revealed by varying development temperature. While this does not exclude the possibility that there are direct thermal effects on the physical interaction between mt-tRNA and tRNA synthetase, it demonstrates that temperature is not required to expose this genetic interaction and suggests a more general physiological explanation – external and internal contexts that place greater demand on energy metabolism expose deleterious effects of genetic interactions that compromise energy supply.

In some insects, including *Drosophila*, pupal eclosion behavior is under control of the circadian clock (Kyriacou *et al.* 1990, Paranjpe *et al.* 2005). Under cyclic or rhythmic photoperiodic regimes, the circadian clock entrains to the light cycle and eclosion behavior is gated such that pupae will delay the initiation of eclosion behavior until lights off in order to synchronize eclosion with dawn. In arrhythmic photoperiodic environments, however, eclosion behavior is unregulated and pupae initiate eclosion behavior as quickly as possible. While compatible mito-nuclear genotypes developed under arrhythmic photoperiods experienced an unregulated acceleration of development, incompatible mito-nuclear genotypes do not appear to have the energetic capacity to similarly accelerate growth. (*simw^501^*);*OreR* larvae also have significantly reduced thermal plasticity for metabolic rate (i.e., they have a very low *Q_10_*) (Hoekstra *et al.* 2013). Thus under two independent contexts that normally accelerate growth (arrhythmic photoperiod and warm temperatures), this mito-nuclear incompatibility appears to limit larval growth. Mito-nuclear incompatibilities likely limit the scope for growth, potentially because incompatibilities compromise ATP production and result in energy supplies that are very close to demand.

### Ontogeny of the energy budget

Even at temperatures that most exacerbate mito-nuclear effects on larval metabolic rate and survivorship (Hoekstra *et al.* 2013), we found that the effects of this mito-nuclear incompatibility on adult metabolic rate are minimal. This suggests that the deleterious effects of mito-nuclear incompatibility on metabolic rate and survivorship may be alleviated by the cessation of growth. Larval growth in *D. melanogaster* proceeds extremely quickly and challenges metabolic processes (Church and Robertson 1966, Tennessen *et al.* 2011). Furthermore, metabolism during growth is estimated to be 40% to 79% above that of fully developed conspecifics (Parry 1983), and cessation of growth in holometabolous insects is correlated with an ontogenetic decrease in metabolic rate per unit mass (Glazier 2005, Callier and Nijhout 2012, Greenlee *et al.* 2014, Maino and Kearney 2014). Thus, the cessation of growth in *Drosophila* likely results in the excess metabolic capacity needed to compensate adult metabolic rate of incompatible, mito- nuclear genotypes. Hoekstra *et al.* (2013) also observed that for those individuals that survive to pupation, there is no further effect of the mito-nuclear incompatibility on survival through metamorphosis, when metabolic rates decrease to a minimum (Dobzhansky & Poulson 1935, Merkey *et al.* 2011) and those individuals that have committed to pupation appear to have the energy stores needed for successful metamorphosis. Consistent with this, we observed that adult (*simw^501^*);*OreR* attain similar mass at eclosion as do adults with compatible mitochondrial and nuclear genomes. Changes in the energy budget of holometabolous insects across development (e.g., Merkey *et al.* 2011) likely generate important ontogenetic contingency for the fitness effects of mutations in populations. The fate of conditionally neutral alleles, such as those underlying this mito-nuclear incompatibility, will then depend not only on what environments are experienced, but when those environments are experienced in an organism’s lifespan (*sensu* Diggle, 1994).

### Sex-specific costs of reproduction

The small effect that this mito-nuclear interaction did have on adult metabolic rate was sex specific. (*simw^501^*);*OreR* females had significantly depressed metabolic rates when measured at cool temperatures. There was some indication that this was driven by disruption of metabolic plasticity, similar to what we have observed in larvae of this genotype, although of much weaker effect; (*simw^501^*);*OreR* females developed at one temperature and measured at the other temperature had the lowest mass-corrected metabolic rates relative to all other compatible genotypes.

If energy stores in mito-nuclear incompatible individuals are limited, then the maintenance of metabolic rate using an inefficient OXPHOS system might support adult, but at a cost to more energetically demanding function, such as reproduction. We observed this pattern in females, but not in males. Females with incompatible mito-nuclear genomes laid far fewer eggs, while the fertility of males with the same mito-nuclear combination was unaffected. The fecundity defects in (*simw^501^*);*OreR* females are also strongly temperature dependent and involve defects in the development and maintenance of the ovary (Zhang *et al.* 2017). Furthermore, mothers of this genotype developed at 28mC produce defective eggs that have a lower probability of being fertilized and, when fertilized, die during embryogenesis presumably due to insufficient maternal provisioning or the inheritance of sub-functional mitochondria (Zhang *et al.* 2017).

Although we measured only one aspect of reproductive fitness for each sex, this pattern is consistent with a higher cost of reproduction in females (Bateman 1948) and with empirical estimates of the costs of gamete production that suggest that high costs of egg production may specifically constrain female gamete production (Hayward and Gillooly 2011). Oogenesis in *Drosophila* is regulated in response to nutrient availability (Drummond-Barbosa & Spradling 2001), but nutrient checkpoints for spermatogenesis are less well studied (but see e.g., McLeod *et al.* 2010, Yang & Yamashita 2015). Further experiments are warranted to determine whether there are more subtle fertility defects in (*simw^501^*);*OreR* males, as cytoplasmic effects on sperm morphology and viability have been measured in seed beetles (Dowling *et al.* 2007b). However, these viability differences do not contribute to cytoplasmic effects on sperm competition in seed beetles or in *D. melanogaster* (Dowling *et al.* 2007c, Friberg & Dowling 2008). Our findings suggest that homeostasis for metabolic rate (or ATP production) combined with potentially differential costs of gametogenesis, may generate sex-specific allocation tradeoffs between maintenance and reproduction as a consequence of genetic variation in metabolic processes.

## Conclusion

Our observation that the degree of expression of a mito-nuclear incompatibility correlates with energetic demand – among developmental treatments that accelerate growth rate, across developmental stages with substantial differences in the cost of growth, and between sexes with putatively different costs of reproduction – suggests that the phenotypic effects of genetic interactions that impact metabolism will depend broadly on the context of energy use and the metabolic cost of producing focal traits. These energetic contexts may change as a function of the external (e.g. temperature or photoperiod) or internal (e.g., life stage, sex or tissue) environment in which a trait is expressed. The context-dependent genotype-phenotype relationships that we describe exemplify how energy allocation principles and design constraints can generate complicated environment-dependence, potentially confounding attempts to define fitness in energetic terms (e.g., Bruning *et al.* 2013). Yet, many components of fitness are expected to depend on the pathways of metabolism, and the variable phenotypic expression of mutations in these pathways presents a dynamic and perhaps challenging context for both purifying and adaptive selection, as context-dependent mutational effects only experience selection in a fraction of possible environments (Van Dyken and Wade 2010). This weakening of selection may be an important contribution to the maintenance of genetic variation for metabolism and life-history traits in populations.

## ACKNOWLEDGEMENTS

The authors thank members of the K. Montooth, C. Meiklejohn and J. Storz labs for feedback on this manuscript, and acknowledge funding support from NSF IOS Award 1149178 to KLM.

## AUTHOR CONTRIBUTIONS

LAH and KLM conceived and designed the study, LAH, CRJ, KMM and KLM collected and analyzed the data, LAH and KLM drafted the initial version of the manuscript, and all authors contributed to later versions of the manuscript.

## DATA ACCESSIBILITY

All data will be available in Dryad upon publication of the manuscript.

